# The small genomics lab experience optimizing data cold storage

**DOI:** 10.1101/2025.03.18.643355

**Authors:** Elisha D. O. Roberson

**Affiliations:** Washington University in St. Louis, Departments of Medicine & Genetics, Division of Rheumatology, St. Louis, MO 63110

## Abstract

Translational research is often a collaborative enterprise that involves basic science researchers, clinicians, and experts in genomics and bioinformatics. While there are central university and industry cores to support data generation, long-term storage often falls to the individual investigators. We frequently fulfill the role of long-term FASTQ file storage for our collaborators. To reduce our cold storage space, we tested the space savings for gzip and zstandard algorithms on an old set of FASTQ files. We found that zstandard had a better overall compression ratio than the best gzip algorithm, amounting to more than 20% space savings overall compared to gzip. It may be worth transitioning to zstandard compression for small, collaborative genomics labs to minimize cold storage costs.

## A. Introduction

The size of datasets produced by research projects has increased in number and size and diversified in content over time. In microscopy, the resolution, number of images, and ability to take live imaging movies produce large amounts of imaging data. For flow and mass cytometry, cells can be assayed at higher throughput and with multiple markers, resulting in sizable Flow Cytometry Standard files. In genomics, the advent of high-throughput sequencing methods has dramatically reduced the cost of sequencing while increasing the accessibility of the technology to more people.

When most scientifically generated data was analog, long-term storage could be as simple as pasting physical images of gels, organisms, or cells into a binder. Redundant backups would have been challenging since it would require manually copying all necessary data and keeping a record elsewhere. Digital data generation and storage presents both opportunities and challenges. Data storage can be far more compact. What may have taken a whole room of binders to store might fit on one flash drive. Keeping multiple copies of digital data can require less individual effort and is more amenable to automation.

It is also worth noting that while the throughput and resolution of many data types have increased over time, it may often be overly expensive, time-consuming, or impossible to recreate some data points.

Recreating human genomic data requires access to a viable sample of the same tissue taken at the previous time points. These limitations make it critically important to ensure the long-term durability of precious data. Data retention allows for the reproduction of original analyses, reuse in the future, and maximizes return on investment.

A problem with the digital nature of data is that it may be difficult for scientists who aren’t trained in computer science to navigate long-term storage. A side effect of the increasingly digital nature of scientific assays is that labs without computational expertise often collaborate with computational labs for data analysis and interpretation. As a result, many academic units have one or more labs experienced in computation that help the labs around them analyze, store, and maintain high-throughput assay data. We have collaborated with many labs over the years, including work as a core supporting a P30. We preserve less data than a university microscopy or sequencing core but more than most standard labs. We also have a more direct role in long-term data preservation. Herein, I describe our experience migrating archival FASTQ data from gzip to zstandard compression.

## B. Results

### B.1. Identifying raw data and strategies for storage

#### What is raw data?

A critical first question for any group grappling with long-term storage is what constitutes raw data. The answer will, of course, depend on the type of experiments they perform. Our group’s most common data type is high-throughput Illumina sequencing. We consider an unaligned FASTQ file (discussion of the format reviewed here ^1^) without trimming or any other read modification as the right kind of raw data for long-term storage. For imaging, raw data might be a captured image or video without color enhancement, cropping, saturation adjustment, or other modifications.

Most large data files are stored compressed. For raw data, consider whether the compression is lossy (the compressed file is smaller because some data are removed) or lossless (the compressed file is smaller because the data are represented differently). Raw data should be stored in a lossless format to retain the original signals.

#### How important is the data?

Information that is cheaply produced, low volume, and is not unique doesn’t require an extensive long-term storage strategy. The cost of storage might outweigh simply generating the data again. However, if the data were costly or required a rare/unique reagent, it is well worth preserving the information. There are also data retention requirements for grant contracts and publications.

#### Strategies for storage

How does one go about saving data that is precious? People sometimes fall back on using a single machine with storage in a RAID (Redundant Array of Independent Disks) configuration ^2^. RAID does have advantages, primarily allowing for multiple disks to appear as a single mount point, automatic copying of information to multiple drives at once (mirroring), and the ability to reconstruct data if one or more disks fail (striping with parity). The strategy does reduce the probability of data loss from catastrophic disk failure of one (or more) drives. However, the machine is still a single point of failure if there is a building fire.

Instead, data should be stored using a 3-2-1 strategy: three separate copies using two different storage methods and at least one copy is stored offsite. This strategy gives several layers of redundancy. It is far less likely that the data will be lost to a random drive failure, as the probability of three simultaneous random failures is low. Two different storage methods help to ensure that if one storage method (such as spinning disk drives) becomes unsupported in the future, the second storage media (burned disk, flash drive, solid state drive, tape) would still be recoverable. How easy is it to recover data from zip disks now compared to the early 2000s? Technologies can and do become obsolete. The offsite backup is to cover you in case of a disaster.

The definition of “offsite” can vary depending on your desire to ensure the data are recoverable. To ensure that the data are safe from a localized disaster (a building fire, flooding, etc), having a copy elsewhere on campus or at a nearby building is enough. If you want recoverability from a regional natural disaster (hurricane, tornado outbreak, wildfire, or earthquake), store a copy in another geographic region.

#### Ensuring data durability versus data availability

We’re primarily interested in the long-term durability of our access to data and our ability to restore it in the case of data loss. Durability is separate from making the data available to the public or ensuring high availability. Public data deposition could be depositing data in an archive that gives a data object identifier, such as FigShare or Zenodo. Or deposition into a government-subsidized ecosystem, such as the gene expression omnibus, the Sequence Read Archive (SRA), or the European Nucleotide Archive (ENA) ^3–5^. Data that needs high availability for many individuals at once and in different geographic regions might be better served by storage in a cloud storage provider, such as Google Cloud Storage, Azure storage, or Amazon EC2.

Due to our interest in genomic data from human samples, long-term storage must be compatible with the consent form used to enroll a human subject. NIH requests for data sharing and journal requirements for data sharing cannot override the protections for human subjects. In practical terms, that means that individuals consented with older IRB protocols or that otherwise did not explicitly include public data sharing require controlled data access protocols, such as using dbGAP instead of the SRA or ENA. In some cases, the generation of genomic data may be allowed, but sharing is forbidden.

Understanding data security requirements is crucial for genomics groups that collaborate often. The ethical considerations of sharing human genomic data are totally different from model organism data.

Collaborators who work in model organisms may assume that all human data can be archived in public repositories.

### B.2. A disposability model of processed data

Raw data is, of course, only part of the picture. The analysis of raw data is required to build insights. Processed genomic data could include cleaned FASTQ files, aligned BAM files, variant call files, or count matrices ^6,7^. Given the ever-increasing size of data produced by scientific research, how do we reconcile the already difficult job of preserving raw data with similar or larger files produced during analysis?

The answer probably again lies with data importance and use. A gene expression catalog might need to keep the estimated abundance of known genes for each sample but might not need to have all the aligned BAM files on hand. Other projects may not need to keep the processed files available after initial use.

Our approach to this issue has been to view the raw data as invaluable and the processed data as reproducible. Reproducing manual data analysis requires careful documentation of the steps. Generally, we try avoiding manual steps in favor of automating analysis using Snakemake workflows stored in Git repositories that run the analyses in Docker images ^8^. The analysis code is small compared to the data. Regenerating the analysis only requires us to clone the repository, transfer the raw data, and execute the workflow. Using workflows and docker images allows us to keep only the most analyses. The approach also increases the reproducibility and transparency of our research. Another important consideration is how time-bound and costly an analysis is when deciding how disposable the outputs are. It may be worth keeping an archive of all project analyses if they require fourteen weeks of cluster computing time.

### B.3. Lossless zstandard compression reduces cold storage costs over gzip compression

The most common algorithms for genomics data compression are probably gzip and bgzip (a block implementation of a gzip-like algorithm) ^9^. The gzip program was released in 1992 and it has been used for compressing archives ever since. The zstandard algorithm is a more modern alternative that claims higher compression ratios with faster compression and decompression than gzip algorithm (zlib) ^10^. A disadvantage of this design is that it may require far more RAM and CPU resources than a zip algorithm. Gzip may be a more economical choice for short-lived data since it uses less CPU and memory resources. For long-term cold storage, however, the primary cost would be the storage rather than the initial compression. In that case, zstandard may be a superior choice.

Our lab has worked with many collaborators for more than a decade. Because of this, we have accumulated an archive of FASTQ files generated by other labs. This cold-storage pilot included files we hadn’t accessed in more than 4 years, amounting to 274 FASTQ files. The gzip algorithm reduced files to a median of 15.12% of raw file size ± 6.09% median absolute deviation (mad). Zstandard archives were a median of 13.18% of raw file size ± 2.75% mad (Fig. 1). The uncompressed FASTQ files were a total of 5.83 TB in size, the gzip FASTQ files were 1.35 TB, and the Zstd compressed FASTQ files were 1.07 TB. A larger starting file size predicted a lower compression level for gzip and zstandard (5.89 × 10^−16^). Zstandard compression predicted a significantly higher compression ratio (smaller final file size) than gzip (1.82 × 10^−9^). For our cold storage archive, the zstandard compression reduced the disk space by more than 20% compared to gzip.

**Fig. 1.**
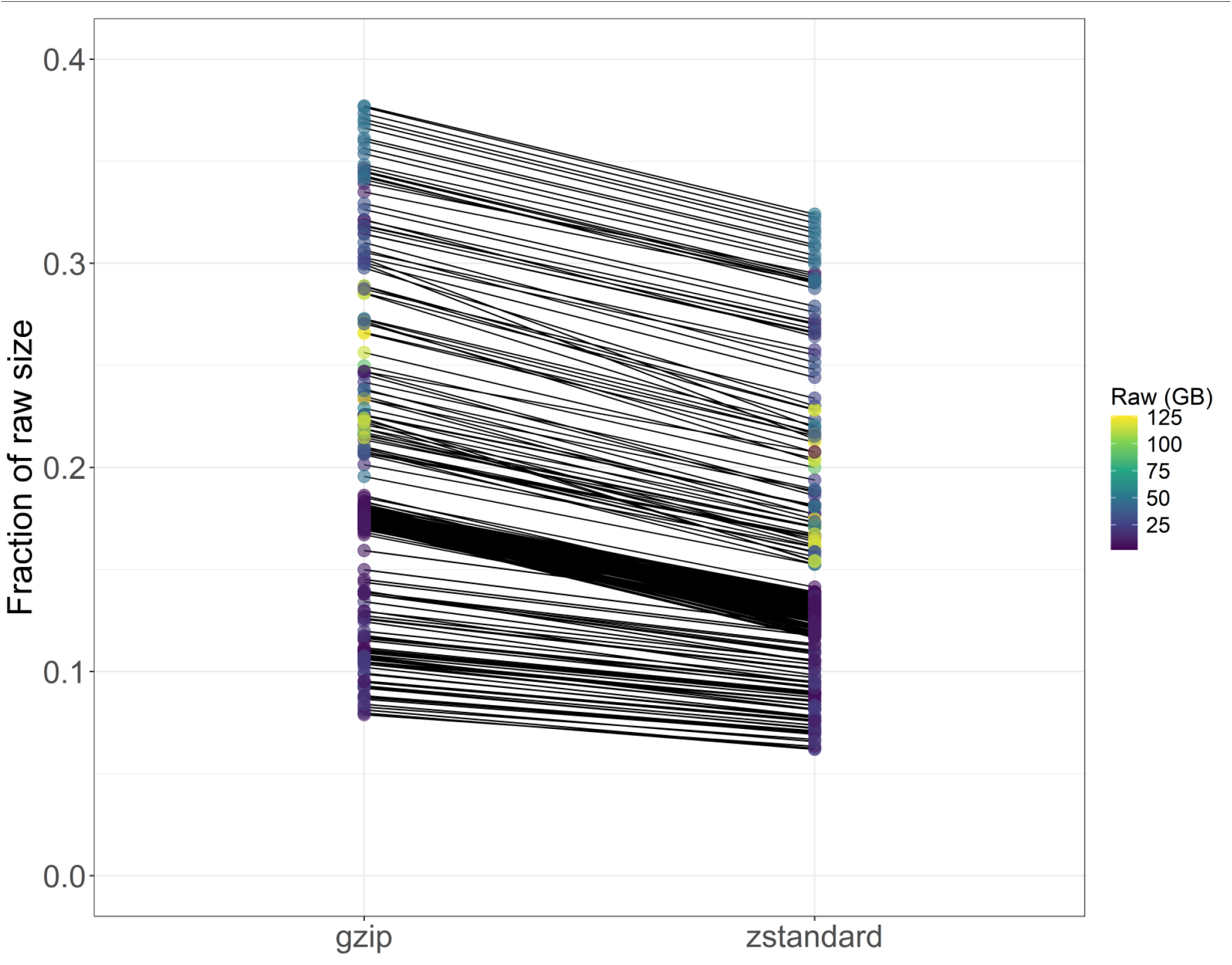
Individual file sizes compared to uncompressed files. Shown are the individual file results for gzip and zstandard compressed files (x-axis). The y-axis is the fraction of the file size compared to no compression. Individual files are connected by lines between compression categories, and the dot color is scaled to the uncompressed file size. In all cases, compressed file sizes were less than half of uncompressed file sizes. In the best cases overall compressed file size was less than 10% of the uncompressed size. Zstandard gave consistently smaller archive sizes.

## C. Methods

We executed the workflow using Snakemake (v6.15.5) plus singularity with Dockerhub image “docker://thatdnaguy/ubuntu_general:v20.04_02”. Each file was decompressed (gunzip v1.10) to raw FASTQ, then compressed with pigz (v2.4) and zstandard (v1.5.5). We calculated the md5 sum (md5sum v8.30) for the raw text file. After zstandard compression, we inflated the file to calculate the md5 sum again. The matching MD5 sums ensured the underlying data were not altered. The pigz compression used the options: --best -p {threads} {input}. The zstandard compression used options: −19 -T{threads} {input}. Both pigz and zstandard used 10 threads. We calculated the byte file size for the raw, gzip, and zstandard files using du (v8.30; du -- block=1 {input} > {output}). We executed the workflow on a Linux server running Ubuntu v18.04.6 LTS that had 64 GB of RAM with an Intel i7-3930K CPU and seven Western Digital Black 6 TB (WD6003FZBX) hard drives mapped as one logical scratch volume.

## D. Discussion

An external perspective of scientific research might be that its function is knowledge generation, spanning discovery on the scale of the entire observable universe to the smallest particles. However, this ideology misses the fact that scientists are people who earn their living by being researchers. The United States biomedical research enterprise is a complex interaction of multiple entities (universities, private industry, and government) that incentivize specific behaviors. Gauging success often relies on poor metrics, including judging research quality by the impact factor of the publishing journal and equating researcher success to citation metrics like h-index or luck in gaining grants. The gamification of scientific success leads to the promotion and retention of those who leverage the metrics to their advantage. Some people try to game the system through overt fraud, leading to an ever-increasing number of paper retractions ^11–13^.

As a field, we’ve incentivized poor proxy metrics instead of research rigor. The increasing number of published papers and retractions makes it crucial to support and incentivize research rigor instead. One part of rigor is maintaining original, unaltered data that underlie key results. Researchers often don’t understand what constitutes original data or how to preserve it for future use. This job may then fall on collaborators with more expertise in informatics.

We have fulfilled this role many times as collaborators in genomics experiments. As a result, we have a cold storage archive of FASTQ files. Storage costs generally decrease over time. In a 3-2-1 strategy, we usually pay for some storage that isn’t on our local machines. In overhauling our archive, we piloted using zstandard compression instead of gzip. Both archive types were substantially smaller than the raw, uncompressed data. Zstandard, however, gave a total cold archive size that was more than 20% smaller than the gzip archives. Savings of that magnitude would scale directly with long-term cold storage costs. For genomics research labs and genomic research collaborators, it may be worth exploring a transition to alternative compression algorithms to save on the associated storage costs.

## Data availability

The file size data is available from FigShare: https://figshare.com/ndownloader/files/53070965

The Snakemake file and analysis code is available from GitHub: https://github.com/RobersonLab/2025-03_FASTQ_compression_paper

## References

1. Cock PJA, Fields CJ, Goto N, Heuer ML, Rice PM. The Sanger FASTQ file format for sequences with quality scores, and the Solexa/Illumina FASTQ variants. Nucleic Acids Res. 2010;38(6):1767–1771. doi:10.1093/nar/gkp1137

2. Patterson DA, Gibson G, Katz RH. A case for redundant arrays of inexpensive disks (RAID). SIGMOD Rec. 1988;17(3):109–116. doi:10.1145/971701.50214

3. Clough E, Barrett T. The Gene Expression Omnibus database. Methods Mol Biol. 2016;1418:93–110. doi:10.1007/978-1-4939-3578-9_5

4. Leinonen R, Sugawara H, Shumway M. The Sequence Read Archive. Nucleic Acids Res. 2011;39(Database issue):D19–D21. doi:10.1093/nar/gkq1019

5. Leinonen R, Akhtar R, Birney E, et al. The European Nucleotide Archive. Nucleic Acids Res. 2011;39(Database issue):D28–D31. doi:10.1093/nar/gkq967

6. Li H, Handsaker B, Wysoker A, et al. The Sequence Alignment/Map format and SAMtools. Bioinformatics. 2009;25(16):2078–2079. doi:10.1093/bioinformatics/btp352

7. Danecek P, Auton A, Abecasis G, et al. The variant call format and VCFtools. Bioinformatics. 2011;27(15):2156–2158. doi:10.1093/bioinformatics/btr330

8. Köster J, Rahmann S. Snakemake-a scalable bioinformatics workflow engine. Bioinformatics. 2018;34(20):3600. doi:10.1093/bioinformatics/bty350

9. Deutsch LP. GZIP File Format Specification Version 4.3. Internet Engineering Task Force; 1996. doi:10.17487/RFC1952

10. Collet Y, Kucherawy M. Zstandard Compression and the Application/Zstd Media Type. Internet Engineering Task Force; 2018. doi:10.17487/RFC8478

11. Cokol M, Ozbay F, Rodriguez-Esteban R. Retraction rates are on the rise. EMBO Rep. 2008;9(1):2. doi:10.1038/sj.embor.7401143

12. Oransky I. Retractions are increasing, but not enough. Nature. 2022;608(7921):9. doi:10.1038/d41586-022-02071-6

13. Else H. Biomedical paper retractions have quadrupled in 20 years - why? Nature. 2024;630(8016):280–281. doi:10.1038/d41586-024-01609-0

